# Emergence of *Acinetobacter baumannii* International Clone 10 predominantly found in the Middle East

**DOI:** 10.1101/2023.10.09.561570

**Authors:** Nabil Karah, Nathan Faille, Frédéric Grenier, Antoine Abou-Fayad, Paul G. Higgins, Leena Al-Hassan, Benjamin A. Evans, Laurent Poirel, Rémy Bonnin, Anette M. Hammerum, Frank Hansen, Rayane Rafei, Monzer Hamze, Xavier Didelot, Santiago Castillo-Ramírez, Simon Lévesque, Sébastien Rodrigue, Bernt Eric Uhlin, Louis-Patrick Haraoui

## Abstract

*Acinetobacter baumannii* is a globally distributed human pathogen. Infections caused by carbapenem-resistant isolates of *A. baumannii* (CRAB) are of great concern, as treatment options are very limited. Despite having among the highest rates reported worldwide, there exists limited genomic data from CRAB strains isolated in the Middle East. Here we report epidemiological, phenotypic, and genome sequencing data (short reads and long reads) on a set of 60 *A. baumannii* isolates belonging to Sequence Type ST158 (Pasteur MLST scheme). They represent a novel international clone (IC), designated IC10, with limited geographic spread beyond the Middle East. Specific antibiotic-resistance genes associated with this clone were identified and data on the plasmid content associated with this lineage are presented.

## Background

*Acinetobacter baumannii* is a Gram-negative bacterium that, over the last four decades, has become one of the most frequently encountered and problematic human opportunistic pathogens, largely due to its antimicrobial resistance profile^1^. As a cause of hospital-acquired infections, *A*. *baumannii* most commonly leads to ventilator-associated pneumonia, wound infections, and catheter-related bloodstream or urinary tract infections. Infections with *A*. *baumannii* are primarily identified in hospitalized patients, with a propensity for those who are immunocompromised or in intensive-care units^2^. It is also reported in trauma victims with secondary infections, notably among civilians and combatants in the context of armed conflicts^3^.

Carbapenem antibiotics, a class of drugs within the large family of beta-lactam antibiotics, have broad-spectrum antibacterial activity. Although the carbapenems imipenem and meropenem exhibited good activity active against *A*. *baumannii* isolated from human infections, nowadays there is a growing population of *A*. *baumannii* encountered in clinical practice are carbapenem-resistant (CRAB). Carbapenem resistance in *A*. *baumannii* often occurs by acquiring an accessory Ambler class D gene, such as *bla*_OXA-23-like_, *bla*_OXA-24/40-like_, *bla*_OXA-58-like_, *bla*_OXA-143-like_, and/or *bla*_OXA-235-like_, or through upregulation of the intrinsic *bla*_OXA-51-like_ gene^4–6^. In addition, several Ambler Class A and B carbapenemases have been reported in *A. baumannii*, including *bla*_NDM_, which most likely originated in an *Acinetobacter* background^7,8^, and particular alleles of *bla*_GES_^9^. Due to the widespread circulation of mobile genetic elements carrying carbapenemase genes among *A. baumannii*, and the subsequent clonal expansion of carbapenem-resistant strains, some regions of the world have CRAB rates exceeding 70%^10^. This worrisome trend has led the World Health Organization (WHO) to list CRAB as one of its three critical priority pathogens for research and development of new antibiotics^11^.

The global clinical population of *A*. *baumannii* is currently dominated by a few epidemic clones, especially international clones (IC) 1 and 2, also known as global clones 1 and 2^12,13^. Other worldwide prevalent clones are IC6 (ST78^Pas^ and ST944^Oxf^) according to the Pasteur Institute (Pas) and Oxford (Oxf) multilocus sequence typing (MLST) schemes respectively, IC7 (CC25^Pas^ and CC229^Oxf^), and IC8 (CC10^Pas^ and CC477^Oxf^) (https://acinetobacterbaumannii.no/overview/global-epidemiology/;^14,15^). IC4 (ST15^Pas^ and ST438^Oxf^) and IC5 (ST79^Pas^ and ST205^Oxf^) have commonly been recognized as major epidemic lineages in Latin America^16^. IC3, corresponding to sequence type (ST) 3 and clonal complex (CC) 928 is rarely encountered nowadays. More recent clones, IC9 (ST85^Pas^) and IC11 (ST164^Pas^), have also been reported^17,18^.

In this article, we present a comprehensive overview of the geographic distribution, epidemiological information, and genomic data of a novel IC of *A*. *baumannii* represented by ST158^Pas^, including the complete genomes of a 60 representative isolates.

## Materials and Methods

### Bacterial isolates

A total of 60 isolates belonging to ST158^Pas^ were included in this study. The isolates were collected between 2007 and 2022 in Kuwait (*n*=28), Egypt (*n*=7), Lebanon (*n*=5), Pakistan and Peru (*n*=4 each), Jordan and Denmark (*n*=3 each), Saudi Arabia (*n*=2), Iraq, Afghanistan, Belgium, and Ireland (*n*=1 each). Four of the Lebanese isolates were cultured from Iraqi (*n*=3) or Syrian (*n*=1) citizens seeking medical care in Lebanon. The three isolates from Denmark had travel histories to Egypt (*n*=2) and Turkey (*n*=1). All the isolates were cultured from human clinical or screening specimens, including blood, respiratory tract, urine, wound, and stool samples.

### Whole genome sequencing

All 60 isolates underwent short-read sequencing, of which 46 were completed by *Université de Sherbrooke* (this study), 3 by co-authors^19–21^ and 11 by previous studies^18,22–30^. For all the 46 *in-house* sequenced isolates, DNA isolation, library construction and genome sequencing were performed according to the manufacturer’s instructions. Following overnight growth at 37°C in lysogenic broth, genomic DNA were extracted using the Quick-DNA Magbead Plus Kit (Zymo Research) and DNA libraries were prepared using the NEBNext Ultra II FS DNA Library Prep Kit (New England Biolabs, NEB). DNA purifications and size selections were made using Ampure XP beads (Beckman Coulter) and quantified using Quant-it PicoGreen dsDNA assay (Thermo Fisher). The quality and size distribution of the DNA was assessed on a Fragment Analyzer using the HS NGS Fragment Kit (Agilent). The pooled samples were then sequenced on a NovaSeq 6000 sequencer (Illumina) as paired-end 250 bp (PE250) read length.

A subset of 17 isolates were selected for long-read sequencing based on the results of their strain typing, genetic resistance patterns, and geographical and chronological distribution. All the long-read sequences were completed at *Université de Sherbrooke*. Extracted genomic DNA was first treated with the NEBNext Ultra II End Repair/dA-Tailing Module (NEB). Then, barcodes from the Native Barcoding Expansion 1-12 & 13-24 from Oxford Nanopore Technologies (ONT) were ligated using the NEBNext Ultra II Ligation Module (NEB). DNA purifications were made using Ampure XP beads (Beckman Coulter). The DNA from different barcoded samples was pooled and the adapter AMII (ONT) was ligated using the NEBNext Ultra II Ligation Module (NEB). Sequencing was done with a R10.4 MinION Flow Cell using a MinION Mk1B (ONT).

### Data Analysis and Bioinformatics

Quality control and trimming of the Illumina reads was done using fastp 0.21.0 with – cut_right –cut_window_size 4 –cut_mean_quality 20 –length_required 30 – detect_adapter_for_pe^31^. Samples with less than 20X coverage were re-sequenced. Unicycler 0.4.9 was used to make assemblies of the trimmed Illumina short reads and ONT long reads when available^32^. Contigs were filtered to retain only those above 500 bp. Taxonomic identification was made on the assemblies using Kraken 2 (2.0.9-beta)^33^.

Two distinct multi-locus sequence typing (MLST) schemes exist for *A*. *baumannii*, known as Oxford and Pasteur schemes^34,35^. The ST of each strain was determined using mlst 2.11 (Seemann T, mlst Github https://github.com/tseemann/mlst), which made use of the PubMLST website (https://pubmlst.org/)^36^. The isolates were also typed using two single-locus sequence typing schemes based on the allelic identity of their *A. baumannii*-intrinsic *ampC* and *bla*_OXA-51-like_ genes^37,38^.

To assess the genetic distance between our isolates and all the other 10 known ICs, we first generated allelic profiles using chewBBACA AlleleCall 2.8.5^39^, and the *Acinetobacter baumannii* core genome MLST (cgMLST) schema from https://www.cgmlst.org. A minimum spanning tree based on the allelic profiles was rendered with GrapeTree 2.2 using the MSTreeV2 method^40^.

Antibiotic resistance genes were found using ResFinder 4.1^41^. Detected *bla*_OXA_ and *bla*_GES_ variants were curated using the Beta-Lactamase database (BLDB)^42^. The presence and nucleotide sequence of resistance genes were manually verified using the QIAGEN CLC genomics workbench 21.0.3 (QIAGEN Digital Insights, https://digitalinsights.qiagen.com/). The presence of neighboring Insertion Sequence (IS) elements was detected using the ISfinder online application^43^. The existence of 24 *A. baumannii* plasmid-borne replicase genes was investigated *in-silico* according to the *A. baumannii* PCR-based replicon typing (AB-PBRT) scheme and relevant studies^44,45^. Plasmids and surroundings of the resistance genes were annotated based on similarities to the GenBank records. Search for similarities was performed using the Basic Local Alignment Search Tool (http://blast.ncbi.nlm.nih.gov/Blast.cgi)^46^. SnapGene Version 5.3.2 (https://www.snapgene.com/) was used to create, visualize, and compare circular maps of the detected plasmids.

Various bioinformatics tools were used to assess the relationships among the isolates and to determine the evolutionary rate of ST158^Pas^. In the initial step, Snippy 4.6.0 was employed to pinpoint the variants as compared to the reference strain Ab-IC10-Kuwait-1, utilizing the assemblies as input (https://github.com/tseemann/snippy). Following this, the outputs generated were utilized in snippy-core to create a full alignment file. This file was then refined further using snippy-clean_full_aln to obtain a clean full alignment output. Subsequently, this output underwent processing through Gubbins 3.0.0, with the parameter –bootstrap 100, to identify recombination events^47^. To isolate conserved sites from the filtered polymorphic site alignment, we employed the snp-sites tool from Snippy.

### Phylogeny and molecular dating analysis

Construction of a Maximum Likelihood phylogenetic tree was achieved by leveraging the capabilities of RAxML 8.2.12, invoking the parameters -N autoFC -f a -B 0.03 -k -m GTRCAT -x 4743 -p 4743^48^. For temporal analysis, the resultant tree was analyzed using the BactDating R package, running for a million iterations^49^. Five *A. baumannii* IC2 strains were used as outgroup for the phylogeny (SAMN02603140, SAMN10716696, SAMEA104305264, SAMN02581277, SAMN13066390). The steps leading to the dated tree were done using 52 samples deemed to have better assemblies (having less than 100 contigs longer than 500 bp).

### Data Availability

Sequencing reads are deposited in GenBank under BioProject number PRJNA1015678.

## Results and Discussion

We identified a total of 110 ST158^Pas^ isolates in the literature by querying the PubMed® database for biomedical literature (https://pubmed.ncbi.nlm.nih.gov/)^18–30^, from co-authors’ strain collections, and/or from the PubMLST (https://pubmlst.org/organisms/acinetobacter-baumannii) and GenBank records. For isolates for which the year of isolation was known, the oldest dated back to 2007 and the most recent to 2022. A large number of the isolates (*n*=100) were collected in or linked to the Middle East (Kuwait, Iraq, Egypt, Saudi Arabia, Lebanon, Jordan, and Turkey) or nearby countries (Tunisia, Afghanistan, and Pakistan). Ten isolates came from Peru (*n*=4, 2012-2016), China (*n*=2, date of isolation not reported), Belgium (*n*=1, 2014), Russia (*n*=1, 2015), Ireland (*n*=1, 2016), and Indonesia (*n*=1, 2022) without a reported history of international travel or recounted linkage to the Middle East. It is worth noting that no ST158^Pas^ isolates have been recovered from Palestine or Israel so far, despite this region struggling with CRAB since the 1990s (unpublished data of 546 Israeli CRAB genomes spanning 1997-2016 sequenced by Louis-Patrick Haraoui). A connection to a military healthcare base was found for 13 isolates, including the 4 Peruvian isolates. The distribution between countries was skewed by the occurrence of at least two nosocomial outbreaks, accounting for 30 isolates from Kuwait^50^ and 7 from Tunisia^51^, or by repeated isolation from the same patient, as for the 3 Jordanian isolates (this study).

We compiled whole-genome sequencing data for 60 of the ST158^Pas^ isolates. Of these, 17 isolates had combined short-and long-read sequencing data, which allowed 12 isolates to have complete assemblies (fully circularized). Using the Oxford MLST system, 57 isolates belonged to ST499^Oxf^ and 3 to ST1717^Oxf^. The latter were all identified in Egypt. ST499^Oxf^ and ST1717^Oxf^ are single locus variants to each other. The difference was related to only one nucleotide change in their *rpoD* locus.

Based on cgMLST comparisons, our ST158^Pas^ isolates did not cluster within any of the other known ICs (Figure 1). Given its wide geographical spread throughout North Africa, Asia, Europe, and South America, ST158^Pas^ was considered as the first ST of a newly designated international clone, termed IC10. All the 60 analyzed IC10 isolates carried the *ampC-43* allele (100% nucleotide identity), which encoded the *Acinetobacter*-derived cephalosporinase (ADC)-117 variant^37^. In all but one of the 60 isolates, the intrinsic *bla*_OXA-51-like_ gene corresponded to *bla*_OXA-65_ (100% amino acid identity). In the remaining isolate, Ab-IC10-Peru-2, the encoded OXA-51-like oxacillinase differed by a single amino acid (T74N) compared to the OXA-65 of the other isolates. As reported by previous studies, the nucleotide sequence of *bla*_OXA-65_ in our isolates showed 3 synonymous nucleotide substitutions in comparison to the first GenBank-deposited allele of OXA-65 (GenBank: NG_049805.1)^25,52^, which was later linked to IC5, corresponding to ST79^Pas^. IC5, which has mostly been found in Latin America, and our proposed IC10 differed substantially by cgMLST (over 2,000 allele differences) as shown in Figure 1. This deviation reveals that the current numbering system of the OXAs is probably well suited for functional comparisons, while it could be misleading in molecular epidemiological studies when based on the DNA sequence^53,54^. Of note, the *bla*_OXA-65_ gene in one of the previously reported ST158^Pas^ (IC10) isolates from Egypt was disrupted by IS*Aba125* (GenBank accession number: JACSTW010000098.1)^52^. Otherwise, no IS element was found in the vicinity of *ampC-43* or *bla*_OXA-65_ among the 60 isolates analyzed in our study.

**Figure 1:**
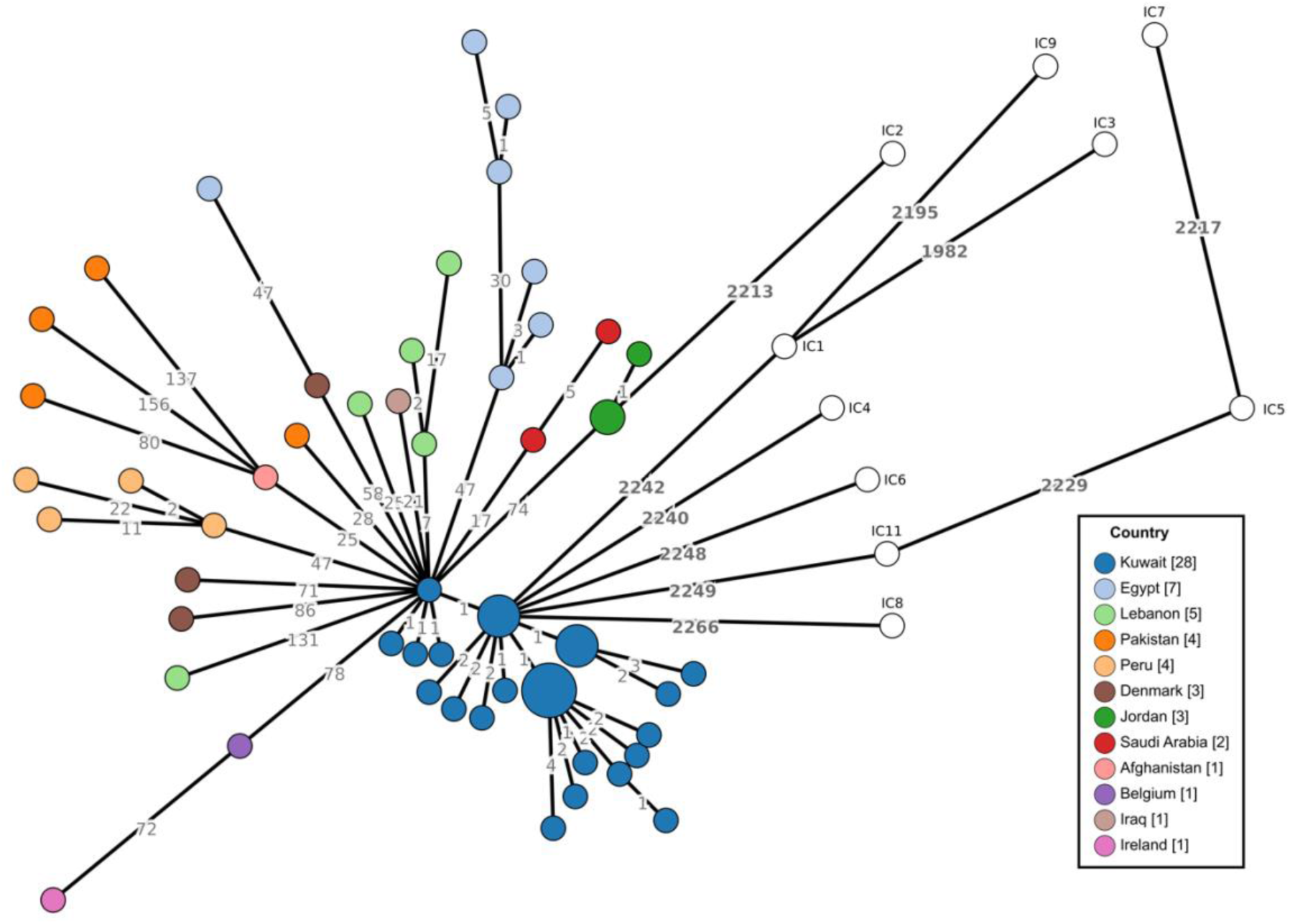
Minimum spanning tree of 60 *A. baumannii* IC10 isolates and 10 International Clones (IC1-IC9, IC11) based on the core genome multi-locus sequence typing (cgMLST) scheme of Higgins *et al.* ^12^. Isolates are color-coded according to the country of isolation and travel history (when applicable). The length of the branches between isolates are proportional to the log transformed number of allelic differences (indicated on the branches) based on the cgMLST scheme. All 10 ICs had at least 2,200 allelic differences with ST158^Pas^/IC10.

### Antimicrobial resistance genes and genetic elements

A total of 23 acquired antimicrobial resistance genes were detected: *bla*_OXA-23_, *bla*_GES-11,-12,-22, and-35_, *bla*_CARB-16 and-49_, *aphA6a*, *aphA1*, *aphA2*, *aacA4*, *aadB* (two different alleles), *strA*, *strB* (two different genes), *cmlA1*, *aadA2b*, *sul1*, *drfA7*, *dfrA1*, *qacE*, *tet(B)*, *tet(X3)*, *msr(E)*, *mph(E)*, *bla*_TEM-1B_, and *aacC3*. The Ambler class D ß-lactamase gene *bla*_OXA-23_ was identified in 52/60 isolates being located within Tn*2008* (48 isolates)^55^ or Tn*2006* (4 isolates, 3 from Jordan and 1 from Denmark with a travel history to Turkey)^56^. In the latter 4 isolates, Tn*2006* was located in *A. baumannii* resistance island AbaR4 that was inserted in the chromosomal *comM* gene, as described in isolates from other ICs^57^.

Five resistance genes (*aacA4*, *drfA7*, *bla*_GES_, *sul1*, and *qacE*) were part of a truncated class 1 integron that was surrounded by two miniature inverted-repeat transposable elements (MITE), forming a mobile element of 7,486 bp^30^. The insertion of this MITE-embarked mobile element, provisionally called MITE*Ab-IC10*, created a characteristic target site duplication of 5-bp^58^. MITE*Ab-IC10* was detected in 51 isolates. In addition, an extended version of this element, including 5 other resistance genes (*aadB*, *cmlA1*, *aadA2b*, *strA*, and *strB*), was detected in 3 isolates (from Lebanon, Ireland, and Belgium). As reported earlier by Mabrouk *et al*., and Valcek *et al*.^28,51^, the extended version of MITE*Ab-IC10* had a length of 13,506 bp (GenBank: KY022424.1 and CP102817.1). Four different *bla*_GES_ alleles were detected, which were *bla*_GES-11_ (49 isolates), *bla*_GES-12_ (2 isolates, 1 from Ireland and 1 from Belgium), *bla*_GES-22_ (1 isolate from Denmark with a travel history to Egypt), and *bla*_GES-35_ (2 isolates, 1 from Egypt and 1 from Denmark with a travel history to Egypt). A previous study reported the occurrence of another allele, *bla*_GES-14_, in seven IC10 (ST499^Oxf^) isolates from Tunisia^51^.

The *aphA6a* gene was detected in 49 isolates, where it was exclusively located on transposon Tn*aphA6*^13^. As reported by other studies^25,28,30,51^, Tn*2008*, MITE*Ab-IC10*, and Tn*aphA6* were all co-located on the same plasmid (see the plasmids section below). The same *aadB* variant (100% nucleotide identity) that was detected in MITE*Ab-IC10* was present in a different genetic context in the chromosome of 2 isolates from Peru. This chromosomal resistance island, around 79,396 bp long and designated IC10-RI, also carried Tn*5* including *aphA2*, *orf* for bleomycin resistance, and *strB* (GenBank: pU00004.1); Tn*5393c* with *strA* and *strB* (GenBank: AF262622.1); IS*CR-2*-related *tetX3*; IS*CR-2*-related *floR*; IS*CR-2*-related *sul2*; part of Tn*7* with *dfrA1*, *sat2*, and *aadA1* (GenBank: KX159450.1 and MN628641.1); *bla*_CARB-16_; and *aadB* with *aphA1*. Some of the resistance parts of IC10-RI were detected on *A. baumannii* strain DT0544C plasmid unnamed1 (GenBank: CP053216.1) while a large part of the background was similar to the genomic records of *Acinetobacter indicus* isolates, such as strain AI18 (GenBank: JAAZRX010000017.1)^59^. IC10-RI’s insertion region was characterized by the occurrence of two conserved 76-bp direct repeats that probably played a role in the acquisition.

In our study, the *tet(X3)* gene was present in 3 isolates that were all collected from Peru. *tet(X3)* was first reported in the genome of isolate *A. baumannii* 34AB, obtained from a pig, where three copies of the *tet(X3)* gene were detected on a large plasmid of around 300,000 bp^60^. A genetic structure related to the one found in p34AB was later reported in *A. baumannii* PW12, an environmental isolate from a Ghanaian Tertiary Hospital^61^. The occurrence of IS*CR2*, Δ*intI*, and *tet(X3)* was shared between p34AB, PW12 and the chromosomal IC10-RI island in our isolates while the region that came directly after *tet(X3)*, which was around 1,100 to 1,200 bp, was completely different. This following region carried *estT*, a serine-dependent macrolide esterase gene^62^. The *estT* gene was present following other *tetX* genes, such as *tet(X5)* and *tet(X6)*^63^. We also detected 3 single changes in the nucleotide identity of *tet(X3)* between p34AB/PW12 and IC10-RI.

Another variant of the *aadB* gene was detected on plasmid pRAY* ^64^ in 6 isolates from Egypt (*n*=3), Saudi Arabia (*n*=2), and Denmark with a travel history from Egypt (*n*=1). The two *aadB* variants shared a nucleotide identity of 98.9% to each other. The macrolide resistance *msr(E)*-*mph(E)* operon was detected in 16 isolates. In 15 of these isolates, the operon was surrounded by two p*dif* sites, creating a genetic element, called module, movable by the XerC–XerD system^65^. The *msr(E)*-*mph(E)* module was carried on a repABSDF_p20001-positive plasmid (6 isolates from Egypt and 2 isolates from Denmark, one with a travel history to Egypt and one to Turkey), a *repAci25*-positive plasmid of 13,351 bp (3 isolates from Pakistan), plasmid of 14,584 bp (3 isolates from Jordan), or plasmid of 8,909 bp (1 isolate from Peru). Each of these four plasmids had its own replicase gene. In contrast, the *msr(E)*-*mph(E)* operon was most likely located on the chromosome, close to an IS*Ec29* element, in one isolate from Denmark with a travel history to Egypt.

The *bla*_CARB-49_ gene, detected in only 2 isolates (Ab-IC10-Belgium-1 and Ab-IC10-Ireland-1), encoded an enzyme differing by 1 amino acid compared to CARB-16. It was surrounded by two copies of IS*Aba1*, forming a 3499 bp composite transposon. We found 2 copies of this transposon, provisionally called Tn*CARB-49*, in the genomic records of Ab-IC10-Belgium-1 (AB32-VUB): one located on its chromosome (GenBank: CP091372.1) while the other was plasmid-mediated (pIC10-1e; GenBank: CP102817.1). We were not able to determine the location of Tn*CARB-49* in the other isolate (Ab-IC10-Ireland-1) since we lacked complete genomic data for this strain.

### Plasmids and other mobile genetic elements

Overall, the isolates carried between 1 and 5 plasmids (Table 1). The most detected plasmid replicase gene was *rep*_ABSDF_p20001_, showing 99.66% nucleotide identity to the original ABSDF_p20001 gene (GenBank: CU468232.1). *rep*_ABSDF_p20001_ was present in all the isolates except for the 4 isolates from Peru. This replicase gene corresponded to Group 12 (GR12) in the Ab plasmid typing scheme^44^. Six different versions of the *rep*_ABSDF_p20001_- positive plasmid, hereafter designated as pIC10-1a to 1g, were detected. The different versions ranged in size from 8,251 bp (pIC10-1f) to 22,905 bp (pIC10-1e). They shared a common backbone of 3,435 bp, including a characteristic region of 4 complete and 1 partial iterons (95 bp). The different pIC10-1 versions carried between 2 and 7 gene modules, each characterized by the presence of two flanking p*dif* sites. Only one of these modules, carried on pIC10-1d, pIC10-1e, and pIC10-1g, was associated with antimicrobial resistance genes, which were *msr(E)* and *mph(E)*, as described above.

The second most common plasmid replicase gene in our isolates was *repAci6*, showing 100% nucleotide identity to the *repAci6* gene of pACICU2 (GenBank: CP031382.1). The corresponding *repAci6*-positive plasmid, designated pIC10-2, was linked to GR6^44^. It had a mean size of around 80,000 bp. pIC10-2 was missing in 1 isolate from Iraq, 1 isolate from Lebanon, and in all the 4 isolates from Peru. Three antimicrobial resistance genetic elements were detected on pIC10-2a, namely MITE*Ab-IC10*, Tn*2008*, and Tn*aphA6*, representing the whole array of resistance genes in most of the host isolates. A variety of versions of this plasmid (pIC10-2a to pIC10-2g) were detected, which were either missing or having an extra genetic element.

The *repAci25* gene was detected only in 3 isolates from Pakistan. As reported elsewhere, *repAci25* had a nucleotide identity of 91% to *repAci4* (GenBank: GU978998.1;^25^). Interestingly, *repAci25* was also carried by plasmid pARA5 hosted by the type strain of *Acinetobacter radioresistens* DSM 6976^T^ (GenBank: AP019745.1). Several plasmids with novel replicase genes were also detected. One gene, encoding the replication initiation protein RepM (WP_019767308.1), was detected in 8 isolates that were all from Egypt or Denmark with a history of import from Egypt. This gene was carried on a plasmid of 5,501 bp (called pIC10-4) that encoded no antibiotic resistance gene. Another *rep* gene was detected in a plasmid of 14,584 bp in the 3 isolates from Jordan. The same plasmid (called pIC10-5) was also carried by an isolate from Saudi Arabia (pMAB25-3; GenBank: CP121593.1). This isolate also belonged to IC10 but was not included in our genomic study since its complete genome was available only recently.

Five other plasmids were detected only in the Peruvian isolates. One plasmid of 105,850 to 113,249 bp (called IC10-7) was present in all 4 isolates from Peru. Another plasmid (pIC10-8, 10,255 to 16,605 bp) was present in 2 isolates. A *repAci9-like* gene was detected on pIC10-6, present in 1 isolate from Peru (Ab-IC10-Peru-2). The *repAci9-like* gene had only 85.5% nucleotide identity to the first reported *repAci9* gene (corresponding to GR8, GenBank: AY541809.1) and should probably be assigned a new number. The backbone of the *repAci9-like*-positive plasmid (8,909 bp, IC10-6) was identical to a plasmid carried by *Acinetobacter lwoffii* strain FDAARGOS (GenBank: CP077374.1). Ab-IC10-Peru-2 carried two other plasmids (pIC10-9 and pIC10-10, 2,858 bp and around 190,000 bp, respectively), with pIC10-10 being very similar to pA297-3 (GenBank: KU744946.1). However, our *in-silico* analysis was not able to determine the exact size and genetic structure of this plasmid or for some of the resistance elements in Ab-IC10-Lebanon-1, - Ireland-1, and -Peru-2.

pIC10-1, the *repAci25*-positive plasmid (pIC10-3), pIC10-5, pIC10-6, and pIC10-8 were all marked by the occurrence of multiple *pdif* sites. Some of the p*dif*-surrounded modules were shuffled between different plasmids. For instance, identical copies of the *orf* (recombinase) module were shared by pIC10-1c, pIC10-3, and pIC10-5. Similarly, identical copies of the *orf* (SMI1/KNR4 family protein) and the *relE*-*yiaG* modules were detected in pIC10-3, pIC10-5, and pIC10-6. However, other modules seemed to have different origins although they had a similar structure as noted for the *orf* (BLUF) module in pIC10-1e and pIC10-3, sharing less than 80% nucleotide identity to each other. Also, the ability of two p*dif* plasmids to merge was indicated by the genetic structures of pIC10-1d and pIC10-1e, although this was not confirmed experimentally.

### Phylogeny and molecular dating analysis

We assessed the phylogenetic relationship of the isolates included in this study. Due to varying sequencing methods and outputs, we performed two ancestral reconstructions: the first containing 52 isolates whose sequencing output included only contigs of 500 bp or more (Figure 2); the other containing all 60 isolates described in this study. Due to concerns about the impact of shorter contigs on the outputs of the ancestral reconstruction, the following results and discussion will focus on the outputs of the first of these two ancestral reconstructions containing 52 isolates.

**Figure 2:**
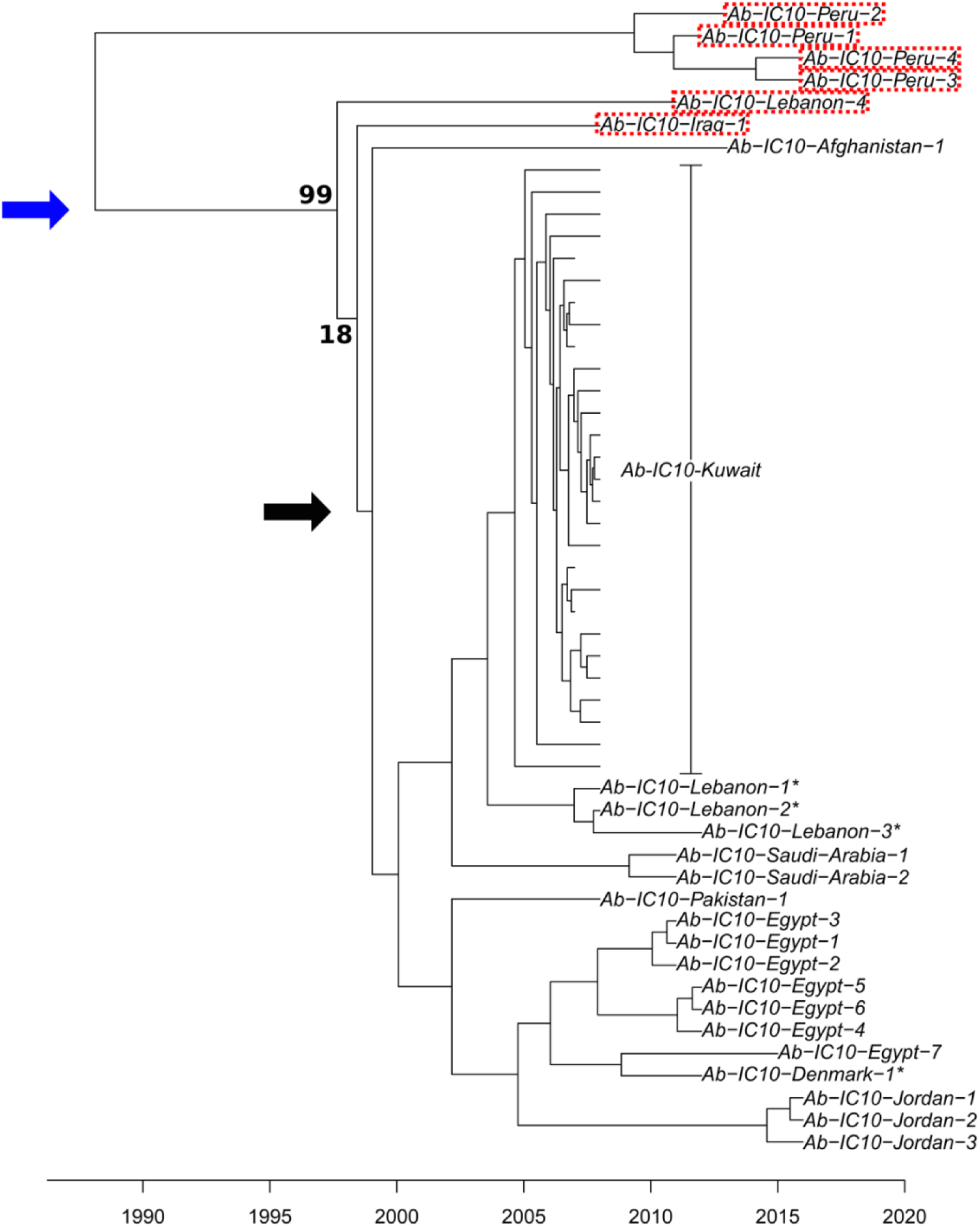
Molecular dating analysis of 52 IC10 isolates. The x-axis indicates the estimated timeline of divergence. The blue arrow indicates the proposed acquisition of pIC10-1, present in all isolates except the 4 from Peru. The black arrow points to the proposed acquisition of pIC10-2. The names of the 6 isolates lacking pIC10-2 are surrounded by red dots. Bold numbers indicate bootstrap values.

Based on the phylogenetic tree of 52 IC10 isolates with high quality sequences, the isolates demonstrated clear grouping by country (Figure 2). One striking finding is the close-to-the-root separation of the 4 isolates from Peru from all the others. Accordingly, the dissemination from/to Peru most likely took place many years prior to our earliest detection of IC10, after which the Peruvian isolates followed a different route of evolution, as denoted by the length of their branch and by their markedly different plasmid content. In addition to containing different combinations of five plasmids not found among any other isolates in this collection (pIC10-6 to pIC10-10), the Peruvian isolates were the only ones lacking pIC10-1.

The phylogenetic tree allowed us to locate a possible single acquisition event of pIC10-1, taking place around 10 years after the split from the Peruvian isolates (Figure 2). The 4 Peruvian isolates also lacked pIC10-2, which was also the case for Ab-IC10-Iraq-1 and Ab-IC10-Lebanon-1. The 6 isolates lacking pIC10-2 appear on branches found on the periphery of the phylogeny. Indeed, the 46 other isolates containing pIC10-2 arose from one branch where a single acquisition event of this plasmid can also be proposed (Figure 2). Following these two single acquisition events, the 46 isolates carrying pIC10-1 and pIC10-2 split between a core group of 45 isolates clustered by country, and the lone isolate from Afghanistan (Ab-IC10-Afghanistan-1).

BactDating^49^ allowed us to estimate the date to the most recent common ancestor (MRCA) of IC10, which was 1987 (with 95% credible interval 1977-1998) with a substitution rate of 1.7 (0.90-2.47) substitutions per genome per year. A possible evolutionary trajectory would be that of an environmental dweller with two notable shifts: 1) transitioning as a human pathogen; 2) acquiring plasmids harboring antibiotic resistance genes facilitating their spread in nosocomial settings and its ability to lead to hospital outbreaks as observed in Kuwait in 2007-8 and in Tunisia 2008-9 .

All isolates contained at least one plasmid. However, given that no single plasmid was shared between the Peruvian isolates and the other Middle Eastern/European isolates in our collection, we present the most likely hypothesis that a strain devoid of any plasmid, was somehow transferred from the Middle Eastern to Peru (Scenario 1), or the other way around (Scenario 2). The Middle Eastern and European descendants acquired sequentially pIC10-1 followed by pIC10-2, while the Peruvian lineage’s descendants went on to acquire different plasmids. Another hypothesis, the separate emergence of two IC10-like clones in the Middle East and in Peru with no link between them, was considered and rejected based on the results of the cgMLST, the phylogeny and the genomic data.

Arguments in favor of Scenario 1 are IC10’s Middle East focus, more specifically its early geographic concentration in several neighboring countries, the timeline outlined above and the historical link between *A. baumannii* and armed conflict-associated infections in the Middle East^66^. It seems possible that IC10 could have emerged as a successful human pathogen in the context of either the Iraq wars or the inter-war period. The latter was marked by a rapidly crumbling healthcare infrastructure in Iraq in the wake of large scale international sanctions^67–69^ affecting all matters of infectious disease practice: access to antibiotics and vaccines; infection control; wound and surgical care; etc. Its spread in the Middle East and beyond can also be linked in certain cases to military operations or bases, such as in Afghanistan and the Naval Medical Research Units in Jordan and Peru. The molecular dating analysis appears to further support Scenario 1 given that the MRCA of the Peruvian isolates appears later than the MRCA of the 48 other isolates. Nevertheless, such an assessment is biased by the limited number of isolates from Peru, so that Scenario 2 cannot be excluded. Access to more IC10 isolates, including some predating the earliest ones from 2007, would further help in establishing which of these two scenarios is correct.

### Conclusions

Fifty-four of 60 IC10 strains we describe were isolated in or related to the Middle East and neighboring countries. At least 13 were linked to military personnel or military healthcare bases. All the IC10 isolates carried specific genetic signatures with regards to their OXA-51-like and ADC enzymes (OXA-65 and ADC-117, respectively), and most carried the *bla*_OXA-23_ carbapenemase gene. Based on their plasmid content, the evolutionary pathway of 4 strains isolated in Peru was strikingly different than that of the 56 other strains we analyzed.

As infections with *A. baumannii* are becoming increasingly more challenging to treat, primarily in low- and middle-income countries, there is a pressing need to expand surveillance of this pathogen. This should be achieved through concerted efforts at typing, sequencing, and strain sharing to facilitate the detection of novel clones, resistance genes and mobile genetic elements. We hope this study and other recent investigations^18,70^ represent the beginning of much-needed efforts to expand our understanding of the growing diversity observed among *A. baumannii* isolates. Together, these contributions will participate in reducing the burden of multidrug-resistant clones by enabling their rapid detection and the deployment of effective infection control measures.

## Notes

### Competing Interest Statement

The authors have declared no competing interest.

